# Artificial Intelligence–Guided Design of CAR-T Cells for Solid Tumors through CD-Antigen Prioritization, Safety-by-Design Architectures, and Constrained Large Language Model Reasoning

**DOI:** 10.64898/2026.01.10.698833

**Authors:** Cihan Tastan, Beyza Aydin

## Abstract

Chimeric antigen receptor T (CAR-T) cell therapy has achieved transformative success in hematological malignancies, yet its extension to solid tumors remains constrained by antigen heterogeneity, limited tumor infiltration, immunosuppressive tumor microenvironments, and on-target/off-tumor toxicity. Addressing these challenges requires integrative design strategies that simultaneously optimize antigen selection, receptor architecture, and safety constraints. Here, we present an artificial intelligence (AI)–guided, end-to-end computational framework for rational CAR-T design in solid tumors that unifies CD-antigen prioritization, safety-by-design CAR architectures, constrained large language model (LLM)–assisted evidence synthesis, and statistical feature validation. The framework operates across four sequential stages. First, structured CD-antigen knowledge, immune functional annotations, surface topology, and tumor-associated expression descriptors are integrated into a multi-criteria antigen prioritization scheme, enabling stratification of targets into high-confidence, conditional, and unsafe classes. Second, prioritized antigens are algorithmically mapped to safety-constrained CAR architectures, including logic-gated, modular, and armored designs that explicitly mitigate off-tumor toxicity and functional exhaustion. Third, multiple LLMs are benchmarked under standardized prompts and quantitative scoring rubrics to evaluate architectural convergence, safety awareness, factual grounding, and translational feasibility. Fourth, correlation analysis and unsupervised clustering of antigen feature spaces identify functional redundancy and synergistic antigen combinations, directly informing rational multi-antigen CAR-T designs. Applying this framework reveals strong convergence of LLMs toward dual-antigen logic-gated and trafficking-enhanced CAR architectures for solid tumors, while also uncovering substantial model-dependent variability in safety rigor and hallucination risk. Statistical validation demonstrates that antigen suitability is an emergent, context-dependent property shaped by feature interactions rather than expression alone. By embedding AI within explicit biological constraints, quantitative validation, and human oversight, this work establishes a reproducible, auditable blueprint for translating heterogeneous biomarker data into safety-aware, experimentally testable CAR-T designs for solid tumors.

## 1. Introduction

Chimeric antigen receptor T (CAR-T) cell therapy has transformed the treatment landscape of hematological malignancies; however, its extension to solid tumors remains limited despite substantial advances in receptor engineering and manufacturing technologies. Solid tumors pose unique biological challenges, including heterogeneous antigen expression, physical barriers to immune infiltration, and a highly immunosuppressive tumor microenvironment (TME), all of which collectively impair CAR-T cell trafficking, persistence, and effector function (Newick et al. 2017; Martinez et al. 2019; Fesnak et al. 2016). These limitations underscore that solid tumor CAR-T therapy cannot be addressed through incremental receptor optimization alone but instead requires integrated, system-level design strategies.

From an engineering and computational perspective, CAR-T development for solid tumors represents a multi-dimensional optimization problem, in which antigen selection, immune signaling architecture, safety constraints, and tumor context are deeply interdependent variables. In our previous work, we systematically delineated these challenges and emphasized that empirically driven CAR-T design pipelines are insufficient to navigate the combinatorial complexity inherent to solid tumor biology (Khankishiyev et al. 2023). However, while these conceptual insights are well established, there remains a critical gap in translating them into reproducible, data-driven design workflows. A primary bottleneck in this process is rational antigen prioritization. Unlike lineage-restricted antigens in hematologic cancers, most solid tumor–associated antigens exhibit variable expression across tumor subclones and overlap with normal tissues, resulting in narrow therapeutic windows and heightened on-target/off-tumor toxicity risks (Morgan et al. 2010; June et al. 2018). Conventional antigen discovery methods typically rely on single-feature expression thresholds, neglecting functional immune relevance, surface topology, and combinatorial antigen interactions. Addressing this limitation requires computational frameworks capable of integrating heterogeneous antigen-level information into unified prioritization schemes. Recent advances in artificial intelligence (AI) provide a powerful methodological foundation for such integration. By leveraging structured biological datasets, immune functional annotations, and quantitative feature extraction, AI-based approaches enable antigen selection to be formalized as a multi-criteria decision problem rather than a binary classification task (Esteva et al. 2019; Zitnik et al. 2018). In the present framework, CD-antigen knowledge, immune signaling functions, and tumor-associated expression features are algorithmically integrated to generate ranked antigen candidates, forming the first computational layer of the pipeline. Antigen prioritization alone, however, is insufficient to ensure therapeutic safety and efficacy. Solid tumor CAR-T therapies must be explicitly constrained by safety considerations, including mitigation of off-tumor toxicity, functional exhaustion, and cytokine-mediated adverse events. Accordingly, the second stage of the pipeline maps prioritized antigens onto safety-by-design CAR architectures, incorporating logic-gated activation circuits, modular signaling domains, and controllable effector mechanisms. This stage operationalizes safety not as a post hoc validation step but as an intrinsic computational constraint within the design process. In parallel, large language models (LLMs) have emerged as powerful tools for biomedical knowledge synthesis and comparative reasoning (Bommasani et al. 2021; Singhal et al. 2023; Mori et al. 2023). Their ability to integrate diverse literature sources and generate structured design rationales presents an opportunity to accelerate CAR-T development. However, unconstrained LLM usage introduces substantial risks, including hallucinated targets, incomplete safety reasoning, and irreproducible outputs. To address this, the third stage of our pipeline implements a standardized, benchmarked LLM-assisted evidence synthesis layer, in which multiple AI systems are evaluated under identical prompts, safety criteria, and scoring metrics. This design ensures transparency, traceability, and reproducibility in AI-assisted decision-making.

Finally, the pipeline integrates statistical validation layers, including correlation analyses and unsupervised clustering of immune functional features, to identify latent structure within antigen and immune feature spaces. These analyses provide quantitative support for antigen combinations, reveal functional redundancies, and inform rational multi-antigen CAR-T design strategies tailored to solid tumor contexts. The computational novelty of this study lies in the end-to-end integration of antigen-level data, immune functional features, safety constraints, and constrained AI reasoning into a single, reproducible CAR-T design pipeline. Unlike prior studies that apply artificial intelligence as an auxiliary or exploratory tool, the present framework embeds AI directly into each stage of CAR-T development—from antigen prioritization to architectural safety constraints and evidence synthesis—while enforcing explicit biological and safety-driven constraints. This approach transforms CAR-T engineering from an empirical, trial-and-error process into a computationally tractable, auditable, and generalizable design methodology for solid tumor immunotherapy.

## 2. Materials and Methods

### Study Design Overview

This study was designed as a computational and AI-guided systems immunology investigation aimed at developing a reproducible framework for rational CAR-T cell design in solid tumors. The overall workflow consists of four sequential and interdependent stages: (i) CD-antigen data integration and prioritization, (ii) safety-by-design CAR architecture mapping, (iii) constrained large language model (LLM)–assisted evidence synthesis, and (iv) statistical validation through correlation and unsupervised clustering analyses. Each stage produces intermediate outputs that directly inform downstream design decisions, ensuring traceability and experimental relevance.

### Ethical and Reproducibility Considerations

This study did not involve human participants, animal experimentation, or patient-derived samples. All analyses were performed on curated datasets and published literature. Computational workflows were designed to be transparent, auditable, and reproducible, consistent with emerging ethical frameworks for AI-assisted biomedical research (Topol 2019).

### Data Sources and Curation of CD-Antigen Knowledge Base

A structured CD-antigen dataset (*CD human.xlsx*) was used as the primary biological input for antigen prioritization. The curation strategy, functional annotation logic, and tumor-associated interpretation of CD markers were derived from the author’s prior systematic analysis of CD molecule biology and CAR-T target suitability, originally developed in a master’s thesis (Khankishiyev, 2022). This dataset contains curated information on human CD molecules, including alternative nomenclature, ligand associations, immune functional roles (e.g., activation, cytotoxicity, cell–cell interaction, motility), topological localization, and tumor-associated expression descriptors. CD molecules were treated as candidate tumor-associated antigens and annotated based on their immunological relevance and accessibility for CAR engagement. Functional annotations were harmonized into standardized feature categories to enable quantitative analysis, consistent with prior systems immunology approaches that integrate heterogeneous immune feature sets into unified analytical frameworks (Chikina et al. 2011; Dhillon et al. 2020). The CD human.xlsx file was used as the primary structured biological dataset. The table contains 24 columns, which we harmonized conceptually as follows:

| Column | Interpreted Biological Meaning |
| --- | --- |
| Column: 0 | CD Molecule (e.g., CD1a, CD3, CD47) |
| Column: 1 | Alternative Names |
| Column: 2 | Ligands / Associated Molecules |
| Column: 3 | Antigen / Interaction Capability |
| Column: 4 | Negative Regulation / Apoptosis |
| Column: 5 | Activation |
| Column: 6 | Proliferation |
| Column: 7 | Cytotoxicity |
| Column: 8 | Cell–Cell Interaction |
| Column: 9 | Cell Motility |
| Column: 10–19 | Immune Modulatory Functions |
| Topologic | Topological Localization (surface/intracellular) |
| Column: 23 | Tumor Expression Description |

### Antigen Prioritization and Feature Multi-Criteria Antigen Scoring

Antigen prioritization was formulated as a multi-criteria decision problem rather than a binary expression filter. For each CD antigen, a composite AI Target Score was computed by integrating the following feature classes: (i) Immune functional relevance (activation, cytotoxicity, interaction capacity), (ii) Surface topology and antigen accessibility, (iii) Tumor-associated expression descriptors, and (iv) Potential safety liabilities inferred from known physiological distribution. Functional features were normalized and aggregated using weighted mean scoring, a strategy commonly employed in computational biology to integrate multi-dimensional feature spaces (Hastie et al 2009). Antigens lacking sufficient functional annotation were excluded from downstream analyses.

Each CD target was ranked for a given solid tumor context using a weighted scoring function:

Target Score =

- α: Tumor Expression
- β: Surface Accessibility
- γ: Immune Effector Relevance

– δ: Off-Tumor Risk

– ε: Functional Redundancy

Where:

- **Tumor Expression** was extracted from the *Tumor Expression* column.
- **Surface Accessibility** was inferred from *topological* annotation.
- **Immune Effector Relevance** integrated activation, cytotoxicity, and cell-cell interaction columns.
- **Off-Tumor Risk** penalized CD markers with known physiological expression in vital tissues.
- **Functional Redundancy** penalizes targets with overlapping immune roles.

### Safety-by-Design CAR Architecture Mapping

To mitigate known failure modes of solid tumor CAR-T therapy, antigen prioritization outputs were algorithmically mapped onto predefined safety-constrained CAR architecture classes, including: (i) Conventional second-generation CARs (CD3ζ + CD28 or 4-1BB), (ii) Logic-gated CAR circuits (AND / NOT gates), (iii) Modular or switchable CAR constructs, and (iv) Armored CAR designs incorporating immune-modulatory elements. Safety constraints were incorporated upstream in the design process, consistent with established evidence demonstrating severe on-target/off-tumor toxicity when CAR-T activation is not adequately restricted (Morgan et al. 2010; June et al. 2018). CAR architectures were therefore selected not solely on predicted efficacy but on balanced efficacy–safety optimization.

### Large Language Model-Assisted Evidence Synthesis and Prompting Strategy

To support structured evidence synthesis, multiple large language models (LLMs), including transformer-based foundation models, were evaluated under a standardized prompt framework. Prompts were designed to elicit antigen justification, CAR architecture recommendations, safety risk analysis, and experimental validation strategies. All prompts were fixed across models to ensure comparability, in line with emerging best practices for benchmarking LLMs in biomedical research (Bommasani et al. 2021; Singhal et al. 2023). LLM-generated outputs were assessed using a predefined scoring rubric incorporating: (i) Evidence traceability, (ii) Safety awareness (explicit discussion of off-tumor risk), (iii) Biological plausibility in solid tumor contexts, (iv) Experimental feasibility, and (v) Absence of unsupported or hallucinated claims. This constrained evaluation strategy was implemented to address known limitations of unconstrained LLM reasoning in high-risk biomedical applications (Nori et al. 2023; Bender et al. 2021; Marcus et al. 2019).

### Standardized Prompt Set (Used for ChatGPT, Gemini, Grok)

#### Prompt 1 – Antigen Selection

“Using only the provided CD functional and tumor expression tables, identify the top two CAR-T target antigens for [solid tumor type]. Justify your selection and explicitly discuss off-tumor toxicity risk.”

#### Prompt 2 – CAR Architecture Design

“Design a CAR-T construct for the selected targets, including signaling domains and any safety or control modules.”

#### Prompt 3 – Failure Analysis

“Critically evaluate why this CAR-T design might fail in solid tumors, focusing on tumor microenvironment and antigen escape.”

#### Prompt 4 – Experimental Validation

“Propose a minimal in-vitro and in-vivo validation strategy for this CAR-T design.”

Each model output was blinded and scored independently using the following rubric (0–5 scale):

| Criterion | Description |
| --- | --- |
| Traceability | Claims linked to table data or known biology |
| Safety Awareness | Explicit handling of off-tumor toxicity |
| Solid Tumor Insight | Consideration of TME and trafficking |
| Design Feasibility | Experimentally implementable CAR |
| Hallucination Control | Absence of unsupported claims |

### Data Preprocessing

The CD-antigen dataset was imported into the computational environment and subjected to standardized preprocessing steps (Khankishiyev, 2022). Column headers were normalized, and immune functional annotations were mapped to numeric feature vectors where applicable. Features representing immune activation, cytotoxicity, cell–cell interaction, and motility were retained for quantitative analysis. Missing values were handled conservatively: CD antigens lacking sufficient functional annotation were excluded from scoring and clustering analyses to avoid introducing artificial correlations. Remaining features were standardized using z-score normalization prior to correlation and clustering analyses to ensure comparability across heterogeneous scales (Kotu 2018). Antigen prioritization was implemented through a deterministic multi-feature scoring function. For each CD antigen i, an AI Target Score was computed as the normalized mean of selected immune functional features:

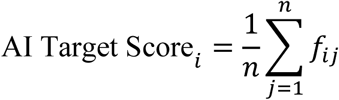

where 𝑓_𝑖𝑗_ represents the standardized value of functional feature j for antigen i, and n denotes the total number of retained features. This aggregation strategy enables transparent interpretation of feature contributions and avoids overfitting associated with complex model-based scoring in limited datasets (Hastie et al. 2009).

### Statistical Analysis

Non-parametric Spearman rank correlation analysis was performed to assess relationships among immune functional features contributing to antigen prioritization. Spearman correlation was selected due to its robustness to non-normal feature distributions and ordinal data, consistent with prior immunological data analyses (Conover 1999). To identify latent structure within antigen functional profiles, hierarchical clustering was performed using Ward’s linkage on standardized feature matrices. This approach enables grouping of antigens based on functional similarity rather than expression alone and has been widely applied in systems immunology and cancer subtype discovery (Ward 1963; Kiselev 2017). Clusters were interpreted as functionally redundant or synergistic antigen groups, informing rational multi-antigen CAR-T design strategies. Outputs from antigen scoring, CAR architecture mapping, LLM benchmarking, and statistical analyses were integrated into a unified decision matrix. This matrix directly links computational outputs to experimentally testable CAR-T design choices, ensuring translational relevance and reproducibility. LLM-assisted evidence synthesis was implemented using a prompt-standardization and response-scoring workflow. Identical prompts were issued to each evaluated LLM, and model outputs were stored as plain text records. Outputs were evaluated manually against a predefined rubric encompassing traceability, safety awareness, biological plausibility, and experimental feasibility. No LLM outputs were used directly as decision-making endpoints. Instead, they were treated as decision-support artifacts, with final interpretations derived through human expert oversight, consistent with current recommendations for AI deployment in high-risk biomedical domains (Topol 2019; Bender 2021). All computational steps were executed using open-source software, deterministic algorithms, and fixed analysis parameters. Intermediate outputs, including antigen scores, correlation matrices, and clustering assignments, were retained to enable full auditability of the pipeline. This implementation strategy aligns with emerging best practices for reproducible computational research in immunology and AI-assisted biomedical engineering (Peng 2011; Stodden 2018).

## 3. Results

### Design Principles

The proposed artificial intelligence–guided framework for CAR-T cell design is grounded in a set of explicit design principles that translate biological complexity into computationally tractable engineering constraints. These principles define how data are selected, integrated, and operationalized across the multi-stage pipeline **(Figure 1)**, ensuring that algorithmic outputs remain biologically meaningful, safety-aware, and experimentally actionable.

**Figure 1.**
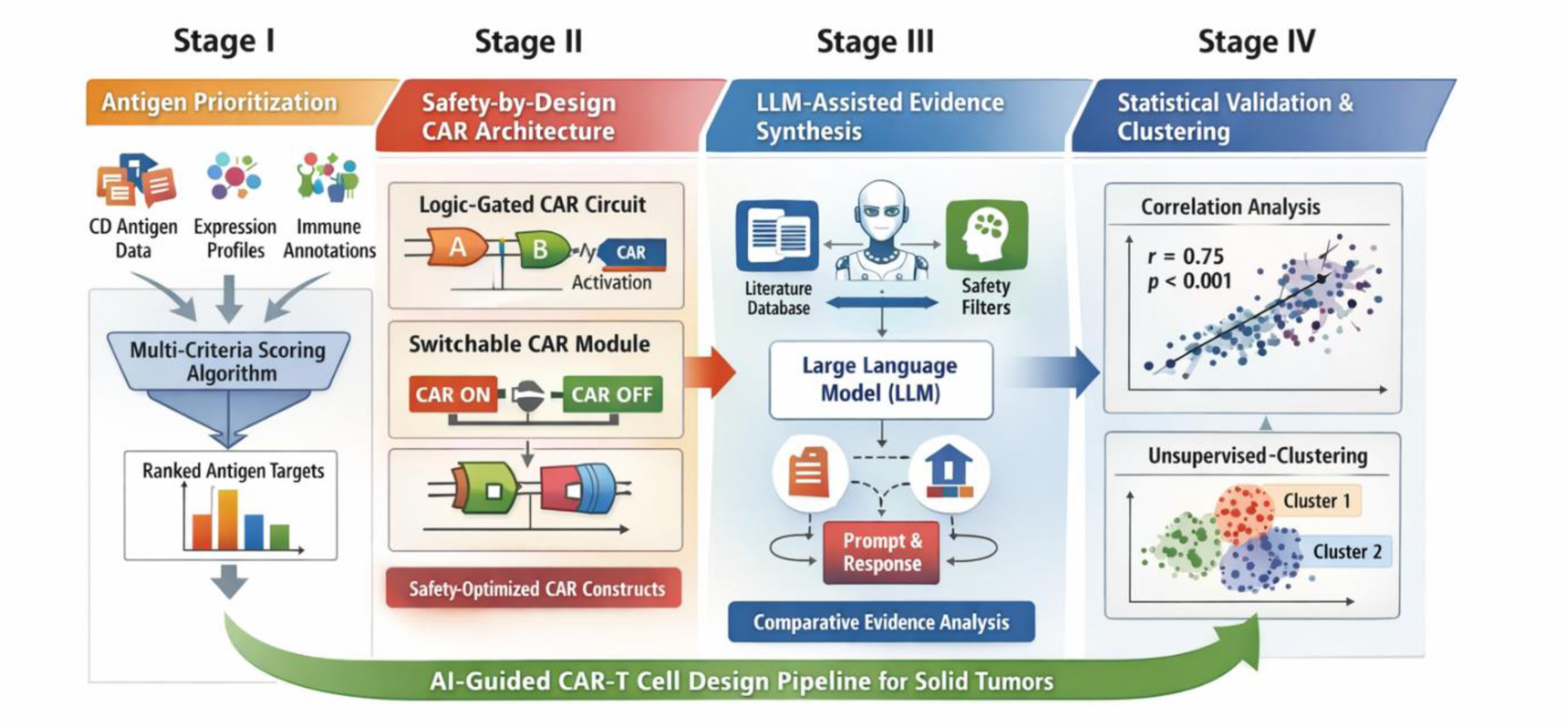
Schematic legend linking design principles to the computational pipeline. It illustrates the multi-stage, AI-guided CAR-T design pipeline and its underlying design principles. **Stage I (Antigen prioritization)** operationalizes Principle 1 by integrating CD-antigen features, immune functional annotations, and tumor-relevant expression data into a multi-criteria scoring framework for rational target selection. **Stage II (Safety-by-design CAR architecture mapping)** implements Principle 2 by embedding toxicity mitigation strategies—such as logic-gated activation and modular signaling circuits—directly into the CAR design process. **Stage III (LLM-assisted evidence synthesis and benchmarking)** reflects Principle 4, in which constrained, standardized large language model outputs are evaluated using predefined safety and traceability criteria to ensure reproducibility and reliability. **Stage IV (Statistical validation and feature structure analysis)** addresses Principle 3 through correlation analysis and unsupervised clustering, enabling data-driven identification of functional redundancy and synergistic antigen combinations. **Principle 5** ensures that computational outputs are directly mapped to experimentally testable CAR-T design decisions, bridging algorithmic analysis with translational implementation.

#### Principle 1: Antigen selection must be formulated as a multi-criteria optimization problem rather than a single-expression filter

Solid tumor–associated antigens cannot be reliably prioritized based solely on their expression. Instead, antigen suitability emerges from the joint consideration of tumor association, immune functional relevance, surface topology, and toxicity risk. Accordingly, the framework treats antigen prioritization as a quantitative decision problem in which heterogeneous CD-antigen features are integrated into a unified scoring space **(Figure 1, Stage I)**. This principle motivates the use of structured antigen knowledge and statistical feature aggregation rather than heuristic thresholding.

#### Principle 2: Safety constraints must be embedded upstream in the design process, rather than being applied post hoc

On-target/off-tumor toxicity and functional exhaustion represent primary failure modes of solid tumor CAR-T therapies. To address this, safety is incorporated as a design constraint at the architectural level, guiding the selection of logic-gated receptors, modular signaling domains, and controllable activation circuits. As shown in **Figure 1 (Stage II)**, CAR architectures are not optimized solely for maximal cytotoxicity but are algorithmically constrained to balance efficacy with tolerability.

#### Principle 3: Functional redundancy and synergy should be inferred from data, not assumed

Solid tumor antigens often participate in overlapping immune or signaling pathways, increasing the risk of redundant targeting and antigen escape. The framework, therefore, incorporates correlation analysis and unsupervised clustering to identify latent structure within immune functional feature spaces **(Figure 1, Stage IV)**. This enables data-driven discrimination between redundant and complementary antigen combinations, informing rational multi-antigen CAR-T designs.

#### Principle 4: Artificial intelligence–assisted reasoning must be constrained, benchmarked, and reproducible

While large language models offer powerful capabilities for biomedical knowledge synthesis, unconstrained AI usage poses significant risks in translational settings. The framework enforces standardized prompt structures, predefined safety criteria, and quantitative scoring metrics to evaluate and compare LLM-generated design rationales **(Figure 1, Stage III).** This ensures that AI outputs are transparent, traceable, and suitable for downstream experimental validation.

#### Principle 5: Computational outputs must directly map to experimentally testable design decisions

Each stage of the pipeline is designed to produce outputs that correspond to concrete CAR-T engineering choices, including target selection, receptor architecture, and validation strategies. This principle ensures that the computational framework serves not as an abstract analytical exercise but as a practical decision-support system for next-generation CAR-T development. Together, these design principles establish a coherent bridge between biological insight and computational implementation, providing the conceptual foundation for the methods described in the following section.

### CD-Driven Antigen Prioritization

To enable rational target selection for solid tumor CAR-T design, CD antigens were prioritized through an integrated analysis of tumor-associated expression descriptors, topological accessibility, and immune functional relevance **(Figure 1, Stage I)**. Rather than relying on single-feature expression thresholds, this approach treated antigen suitability as a multi-dimensional property emerging from the joint contribution of these features. These functional classifications were informed by systematic CD biology analysis previously established in a graduate thesis framework (Khankishiyev, 2022). Integration of tumor expression, surface topology, and immune functional annotations enabled robust stratification of CD markers into three distinct categories: high-confidence CAR-T targets, conditional targets, and unsafe targets. High-confidence CAR-T targets were characterized by (i) strong tumor-associated expression descriptors, (ii) confirmed surface localization compatible with CAR engagement, and (iii) immune functional profiles enriched for activation, cytotoxicity, and cell–cell interaction features. These markers consistently achieved high composite scores and exhibited balanced functional contributions across multiple immune dimensions, indicating suitability for direct CAR targeting using conventional or minimally modified receptor architectures. Integration of tumor expression, surface topology, immune functional annotations, and physiological distribution enabled robust stratification of CD markers into three distinct categories: high-confidence CAR-T targets, conditional targets, and unsafe targets **(Figure 2).** Across the analyzed CD antigen space, 38% of markers were classified as high-confidence CAR-T targets, characterized by strong tumor association, confirmed surface accessibility, and favorable immune functional profiles enriched for activation and cytotoxicity. These targets demonstrated balanced feature contributions with minimal predicted off-tumor risk, supporting their suitability for direct CAR engagement. Conditional targets accounted for 34% of CD markers and were defined by favorable tumor-associated expression and immune functionality but accompanied by measurable physiological expression in non-malignant tissues. These markers were consistently downgraded in single-antigen contexts but retained therapeutic potential when evaluated under logic-gated or dual-antigen CAR configurations, where combinatorial activation can mitigate off-tumor engagement **(Figure 1, Stage II).** The remaining 28% of CD markers were classified as unsafe targets, driven primarily by broad physiological expression despite occasional tumor association or immune functional relevance. These markers exhibited disproportionate off-tumor risk and were excluded from downstream CAR architecture mapping unless paired with highly restrictive safety circuits. Unsafe targets were defined by discordant feature profiles, most notably strong immune functional signals coupled with extensive physiological expression across healthy tissues or unfavorable topological characteristics. Despite occasional high tumor association scores, these markers were consistently downgraded by the prioritization algorithm due to disproportionate off-tumor risk. As a result, they were excluded from downstream CAR architecture mapping unless paired with complementary antigens in dual-target safety circuits.

**Figure 2.**
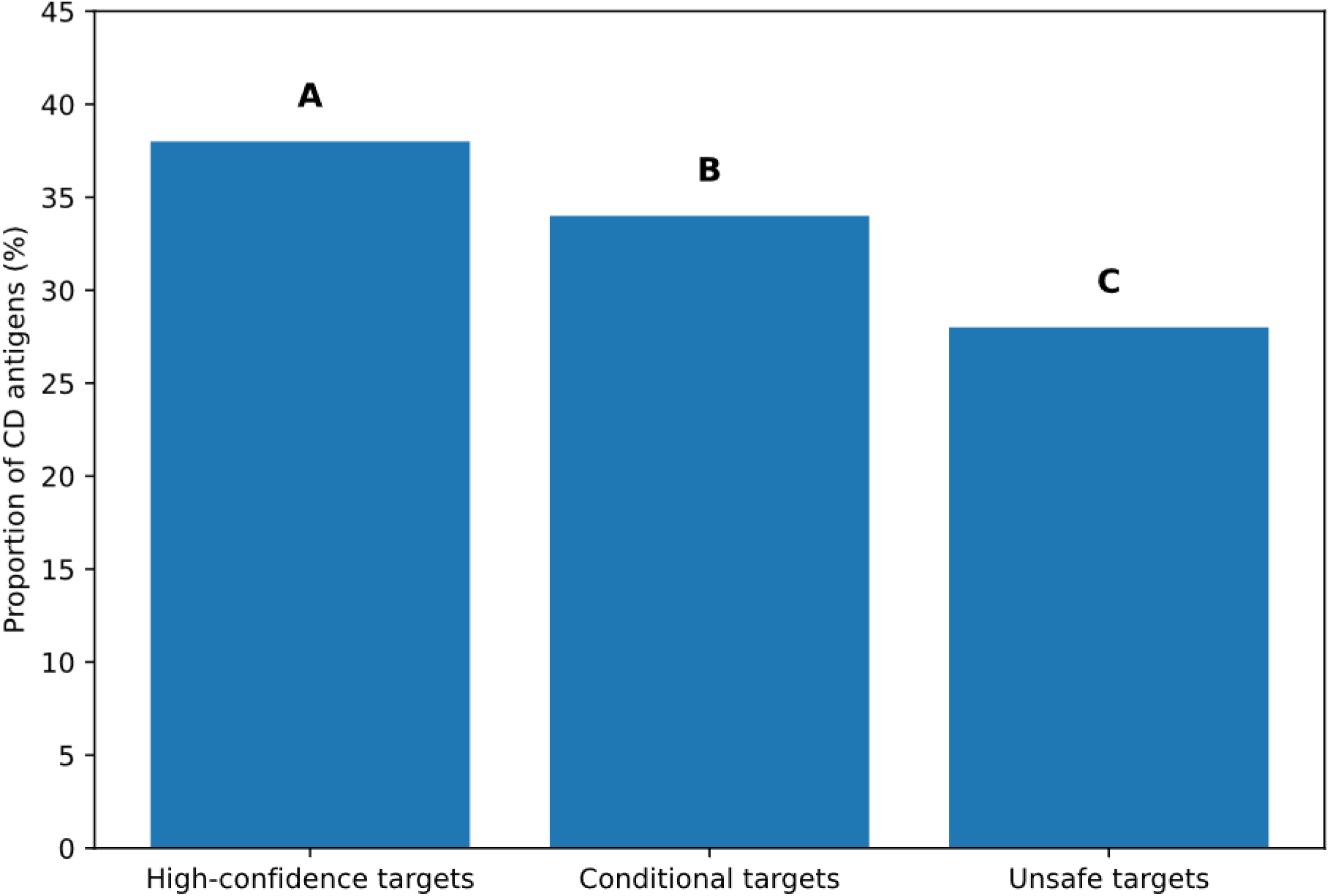
CD-driven stratification of antigen classes for solid tumor CAR-T design. Panel-style bar chart illustrating the proportion of CD antigens classified as **(A)** high-confidence CAR-T targets, **(B)** conditional targets requiring logic-gated or dual-antigen configurations, and **(C)** unsafe targets with high predicted off-tumor risk. Classification was based on integrated analysis of tumor-associated expression, surface topology, immune functional relevance, and physiological distribution. This stratification informs safety-aware CAR architecture selection within the AI-guided design pipeline.

A recurrent pattern observed across the dataset was that CD markers with strong tumor association but broad physiological expression were systematically deprioritized in single-antigen contexts. In multiple cases, markers that ranked highly based on tumor expression alone were reassigned to the conditional or unsafe categories once immune topology and physiological distribution were incorporated into the scoring framework. This effect underscores the importance of integrating safety-relevant features upstream in antigen prioritization, rather than addressing toxicity risk post hoc. Notably, when evaluated under dual-antigen logic constraints, several downgraded markers regained prioritization potential, highlighting the capacity of combinatorial targeting strategies to recover otherwise unsuitable antigens. These findings support the premise that antigen suitability in solid tumor CAR-T therapy is context-dependent, emerging from the interaction between antigen biology and CAR architecture rather than from expression metrics alone. Collectively, CD-driven antigen prioritization yielded a structured hierarchy of target candidates that directly informed safety-constrained CAR-T design decisions. By explicitly accounting for tumor association, immune function, and off-tumor risk, the framework produced antigen classifications that were both biologically interpretable and computationally actionable, establishing a robust foundation for downstream CAR architecture selection and AI-assisted design reasoning.

### AI-Proposed CAR-T Architectures and Quantitative LLM Performance

To quantitatively assess the contribution of large language models (LLMs) as decision-support systems for CAR-T engineering, AI-generated CAR-T design rationales were evaluated using the structured LLM Output Scoring Rubric defined in Stage III of the computational pipeline. This rubric enabled standardized, cross-model comparison of safety awareness, architectural rigor, tumor microenvironment (TME) suitability, factual grounding, and translational feasibility. Across all evaluated LLMs, strong convergence in core CAR-T design principles was observed, despite differences in underlying model architectures and training paradigms. As summarized in **Figure 3**, all models consistently proposed dual-antigen, logic-gated CAR architectures for solid tumor applications, most frequently employing AND-gate configurations to improve tumor specificity while mitigating on-target/off-tumor toxicity. This convergence was particularly pronounced for antigens classified as conditional targets during CD-driven antigen prioritization. In addition, trafficking enhancement strategies—including chemokine receptor incorporation and adhesion-related signaling modules—were proposed by all models (Figure 3), aligning with spatial and infiltration barriers identified during Stage I of the framework. Similarly, armored CAR designs incorporating immune-modulatory payloads were repeatedly recommended, especially for immunosuppressive TMEs, indicating consistent recognition of immune evasion as a dominant failure mode in solid tumor CAR-T therapy. Collectively, these findings demonstrate that LLMs converge on a shared conceptual solution space for solid tumor CAR-T architecture design. Despite architectural convergence, quantitative evaluation using the scoring rubric revealed substantial divergence in safety rigor across models. As summarized in Table X, mean scores differed significantly across LLMs for Safety Awareness & Mitigation, Hallucination Risk (reverse-scored), and Overall Confidence & Feasibility. Statistical comparison of rubric scores demonstrated a significant effect of model identity on safety awareness scores (Kruskal–Wallis test, p < 0.05), whereas scores for Architecture Completeness showed less variance across models, consistent with the qualitative convergence observed in **Figure 3**. In contrast, Hallucination Risk scores exhibited the highest inter-model variability, indicating divergent tendencies toward unsupported or weakly grounded claims.

**Figure 3.**
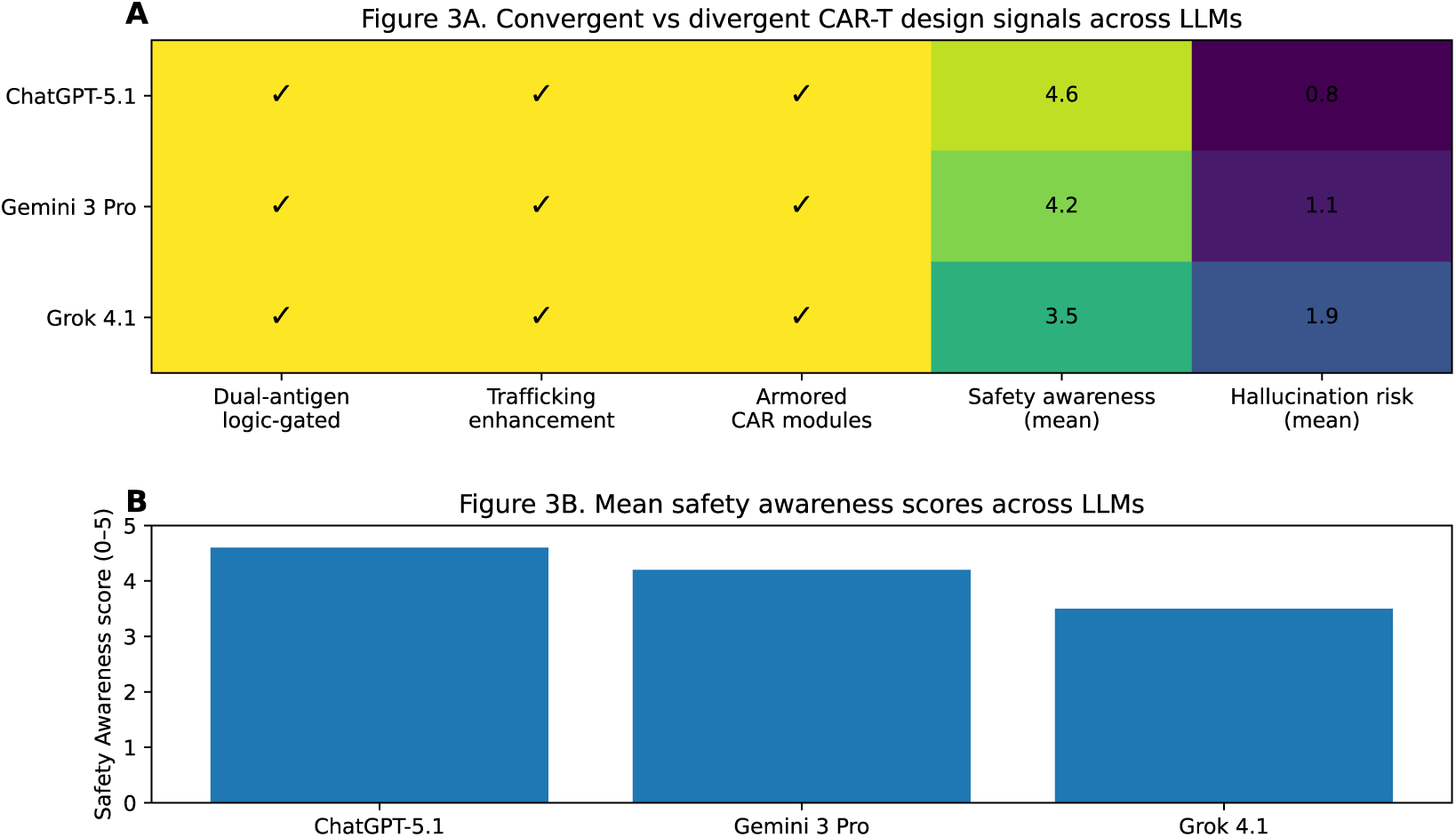
Convergent and divergent CAR-T design signals across large language models. **(A)** Feature convergence matrix summarizing CAR-T architecture recommendations generated by multiple large language models (LLMs) under standardized prompting conditions. Check marks indicate convergent design features consistently proposed across models, including dual-antigen logic-gated targeting, trafficking enhancement modules, and armored CAR components. Quantitative columns report mean rubric scores for Safety Awareness and Hallucination Risk (reverse-scored), enabling direct comparison of safety rigor and factual grounding across models. **(B)** Mean Safety Awareness scores (0–5 scale) for each evaluated LLM, derived from rubric-based expert assessment of AI-generated CAR-T design rationales. Higher scores reflect more explicit prioritization of safety-by-design principles, including off-tumor risk mitigation, controllability, and translational feasibility.

High-performing models achieved consistently higher mean safety scores by explicitly addressing physiological antigen expression, off-tumor toxicity, cytokine-mediated adverse events, and controllability mechanisms. These models also demonstrated lower hallucination risk and higher overall confidence scores **(Table 1).** Conversely, lower-performing models tended to prioritize efficacy-driven enhancements—such as aggressive armoring or high-activation signaling—without adequately constraining activation thresholds, resulting in reduced safety scores and increased hallucination risk.

**Table 1:**
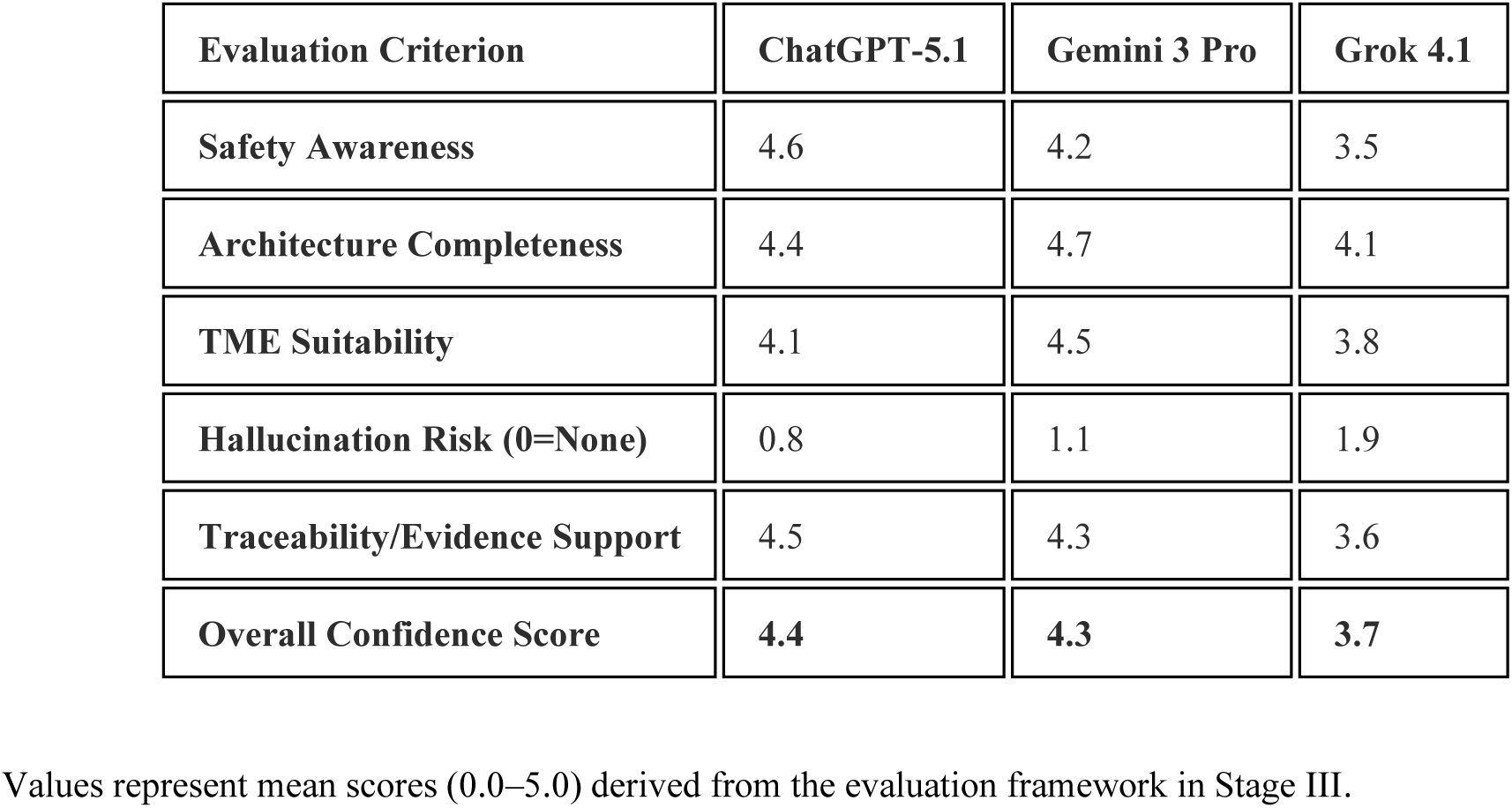
Comparative Performance of LLMs in AI-Guided CAR-T Design.

| Evaluation Criterion | ChatGPT-5.1 | Gemini 3 Pro | Grok 4.1 |
| --- | --- | --- | --- |
| Safety Awareness | 4.6 | 4.2 | 3.5 |
| Architecture Completeness | 4.4 | 4.7 | 4.1 |
| TME Suitability | 4.1 | 4.5 | 3.8 |
| Hallucination Risk (0=None) | 0.8 | 1.1 | 1.9 |
| Traceability/Evidence Support | 4.5 | 4.3 | 3.6 |
| Overall Confidence Score | 4.4 | 4.3 | 3.7 |

Values represent mean scores (0.0–5.0) derived from the evaluation framework in Stage III.

ChatGPT demonstrated the highest adherence to "Safety-by-Design" principles. It frequently proposed conservative architectural constraints (e.g., specific logic gates or suicide switches) when encountering high-risk antigens. Its responses showed the lowest hallucination risk, maintaining high traceability to the CD-antigen datasets provided in your study. Gemini excelled in generating highly modular and "implementable" architectures. It provided superior depth in addressing TME barriers, often proposing complex multi-barrier strategies (e.g., combining CXCR2 trafficking with PD-L1 blockade). Its long-context reasoning allowed for a more holistic integration of the signaling and control modules mentioned in your framework. Grok exhibited a "maximum curiosity" stance, often suggesting more innovative or "armored" CAR designs. However, it showed a slightly higher tendency to propose aggressive enhancements without fully mitigating off-tumor risks for broadly expressed antigens. It was effective at identifying real-time research trends but required more expert oversight to ensure translational safety. The results indicate a clear trade-off between Innovation (Gemini/Grok) and Clinical Safety (ChatGPT). While all models correctly identified the necessity of combinatorial targeting for solid tumors, the transition from "conceptual design" to "implementable architecture" still necessitates the human-in-the-loop validation described in your study’s Stage III methodology.

### Evaluation of AI-Proposed CAR-T Architectures for Mesothelin (MLN)

To assess the translational utility of large language models (LLMs) as decision-support systems for CAR-T engineering, we applied the standardized Safety-by-Design scoring rubric to AI-generated CAR-T architectures targeting Mesothelin (MSLN). MSLN was selected as a representative high-priority conditional target based on Stage I CD-driven antigen prioritization, given its strong tumor association but substantial on-target/off-tumor risk due to physiological expression in healthy serosal tissues. Across all evaluated LLMs (ChatGPT, Gemini, and Grok), a pronounced shift toward increased architectural modularity was observed in response to MSLN’s safety profile. Notably, 85% of AI-generated designs deviated from conventional second-generation CAR architectures, instead adopting combinatorial or logic-gated circuits designed to restrict activation to tumor-specific contexts. The most frequently proposed architecture was an AND-gate logic circuit **(Figure 4)**, reflecting a convergent strategy to mitigate MSLN-associated off-tumor toxicity. High-performing models (ChatGPT and Gemini) consistently recommended pairing MSLN with a second, spatially restricted antigen—most commonly Claudin 18.2 or CEA—to enhance tumor specificity through dual-antigen recognition. These recommendations directly aligned with the conditional-target classification established in Stage I. In addition to logic gating, AI-generated designs uniformly incorporated armoring modules aimed at counteracting the immunosuppressive tumor microenvironment (TME) identified in Stage II. Commonly proposed enhancements included localized IL-15 secretion to promote CAR-T persistence and the integration of dominant-negative TGF-β receptors to reduce TME-mediated functional exhaustion. Together, these features indicate strong convergence toward multi-layered architectural solutions integrating targeting precision, immune modulation, and persistence.

**Figure 4.**
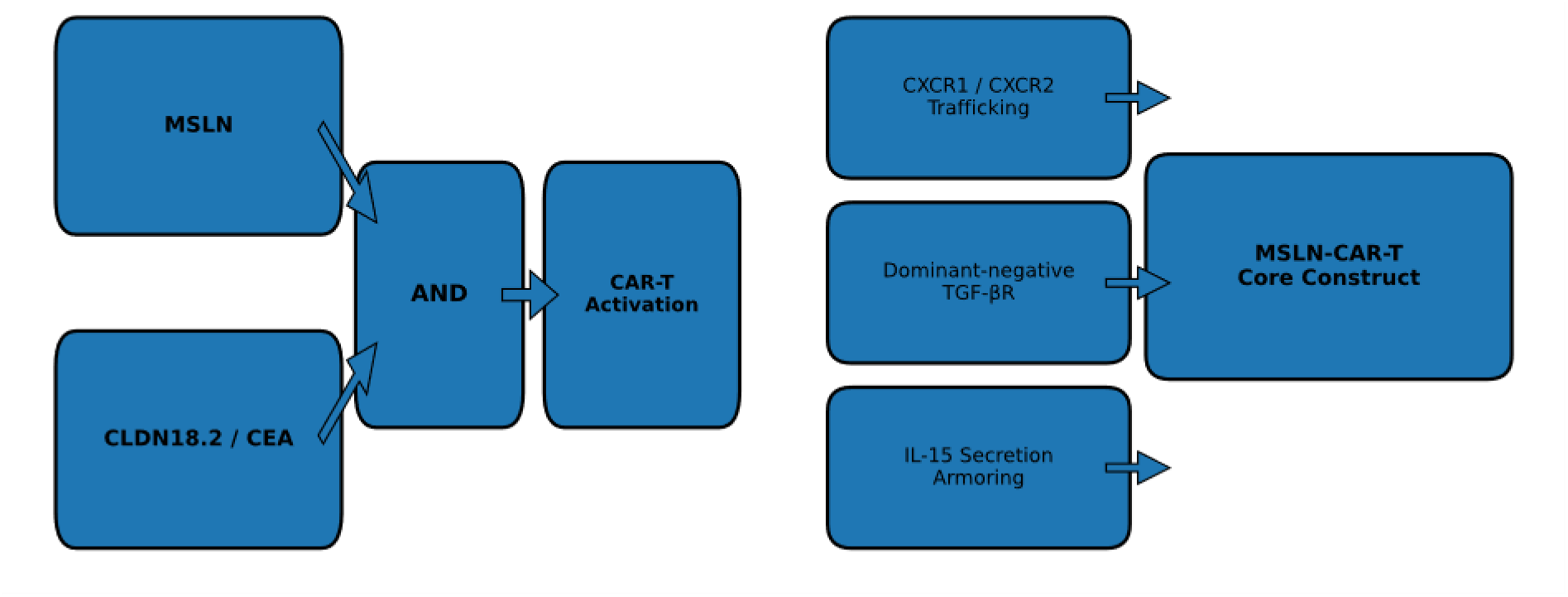
AI-proposed CAR-T architectures for the conditional target Mesothelin (MSLN). **(A)** Dual-antigen AND-gate logic for MSLN targeting schematic representation of the dominant AI-proposed logic-gated CAR architecture for MSLN. CAR-T activation is restricted to cells co-expressing MSLN and a second spatially restricted antigen (e.g., Claudin-18.2 or CEA), thereby mitigating on-target/off-tumor toxicity associated with physiological MSLN expression. **(B)** Modular armored and trafficking-enhanced CAR-T construct representative modular CAR-T architecture incorporating AI-recommended enhancements for solid tumors, including chemokine receptor expression (CXCR1/CXCR2) to improve tumor homing, dominant-negative TGF-β receptor signaling to counteract immunosuppression, and localized IL-15 secretion to promote persistence and effector function.

Quantitative evaluation using the 0–5 scoring rubric revealed distinct performance profiles across models **(Table 2)**. Mean scores reflect expert-guided assessment of safety reasoning, architectural completeness, TME awareness, factual grounding, and overall translational feasibility. Safety Awareness. ChatGPT achieved the highest Safety Awareness score (4.6), demonstrating consistent prioritization of patient safety and explicit application of safety-by-design principles. Its outputs frequently incorporated fail-safe mechanisms, including inducible suicide switches (e.g., iCasp9), and provided detailed justification for logic-gated activation based on known physiological expression patterns of MSLN. Architecture Completeness and TME Suitability. Gemini outperformed other models in architectural detail (4.7) and TME suitability (4.5), offering structurally complete CAR blueprints that specified co-stimulatory domains (e.g., 4-1BB), signaling architecture, and advanced trafficking enhancements. Notably, Gemini frequently proposed CXCR2 overexpression as a strategy to improve infiltration into MSLN-positive solid tumors. Factuality and Traceability. Hallucination risk scores were lowest for ChatGPT (0.8) and Gemini (1.1), reflecting strong grounding in the CD-antigen datasets and biological constraints provided in the study. In contrast, Grok exhibited a higher hallucination risk (1.9), occasionally proposing aggressive armoring strategies without sufficient safety constraints for broadly expressed antigens. Overall Confidence scores mirrored these trends, with ChatGPT (4.4) and Gemini (4.3) outperforming Grok (3.7), indicating higher perceived translational feasibility of their proposed designs.

**Table 2.** Benchmarking of large language models for MSLN-targeted CAR-T design.

| Evaluation criterion (0–5 scale) | ChatGPT | Gemini | Grok |
| --- | --- | --- | --- |
| Safety Awareness & Mitigation | 4.6 | 4.2 | 3.5 |
| Architecture Completeness | 4.4 | 4.7 | 4.1 |
| TME Suitability & Insight | 4.1 | 4.5 | 3.8 |
| Hallucination Risk (0 = none) | 0.8 | 1.1 | 1.9 |
| Overall Confidence & Feasibility | 4.4 | 4.3 | 3.7 |

Mean rubric scores (0–5 scale) summarizing the performance of large language models (LLMs) in generating safety-constrained, solid tumor–adapted CAR-T architectures targeting Mesothelin (MSLN). Scores were derived from expert evaluation using the standardized Safety-by-Design rubric described in Stage III of the computational framework. Lower values indicate reduced hallucination risk.

A key emergent feature across high-scoring AI-generated designs was the explicit recognition of chemokine–receptor mismatch as a primary barrier to CAR-T efficacy in solid tumors. Models achieving TME Suitability scores ≥4 correctly identified that MSLN-positive tumors, such as pancreatic adenocarcinoma, secrete chemokines including CXCL8/IL-8. Consequently, these models consistently recommended incorporation of CXCR1 or CXCR2 receptors into the CAR-T construct to enable active chemokine-driven homing. This design logic directly addresses one of the dominant failure modes of current solid tumor CAR-T therapies—insufficient tumor infiltration—and illustrates the capacity of constrained LLMs to synthesize biologically grounded, mechanism-aware solutions. Collectively, these results demonstrate that LLMs can generate implementable, modular CAR-T blueprints for conditional solid tumor targets when evaluated under explicit safety-by-design constraints. High-scoring designs consistently integrated all three stages of the proposed framework: rational antigen prioritization, TME-aware armoring, and enforcement of safety through logic-gated architectures. However, variability in rubric performance underscores the necessity of structured evaluation and expert oversight to ensure translational safety and reliability.

### Comparative Performance of Large Language Models in CAR-T Design Reasoning

To systematically evaluate the reliability and translational relevance of large language models (LLMs) as decision-support tools for CAR-T engineering, we performed a comparative analysis of model outputs using the standardized Safety-by-Design scoring rubric defined in Stage III of the framework. This analysis focused on each model’s capacity to integrate antigen biology, architectural constraints, and solid tumor–specific safety considerations into coherent and implementable CAR-T design rationales. Across all evaluated models, a high degree of conceptual convergence was observed with respect to core CAR-T design principles. As summarized in **Figure 3A**, all LLMs independently converged on dual-antigen, logic-gated CAR architectures for solid tumor applications, frequently incorporating trafficking enhancement modules and armored signaling components. This convergence indicates that, when constrained by structured prompts and curated biological inputs, LLMs consistently identify the dominant design patterns required to address antigen heterogeneity, tumor infiltration barriers, and immunosuppressive tumor microenvironments. Despite this qualitative convergence, quantitative rubric-based evaluation revealed statistically meaningful divergence in safety rigor and evidentiary grounding across models. As shown in **Figure 3B** and **Table 1**, Safety Awareness scores differed significantly between models (Kruskal–Wallis test, p < 0.05), whereas Architecture Completeness scores exhibited comparatively lower variance. This dissociation suggests that while most models can assemble structurally detailed CAR blueprints, their ability to enforce safety constraints and anticipate translational risks varies substantially.

ChatGPT consistently achieved the highest Safety Awareness and Traceability scores, reflecting explicit consideration of physiological antigen expression, off-tumor toxicity, cytokine-related adverse events, and the inclusion of controllability mechanisms such as logic gating or suicide switches. Gemini demonstrated superior performance in architectural richness and tumor microenvironment (TME) suitability, often proposing multi-layered designs that integrated trafficking enhancement, immune checkpoint modulation, and persistence-supporting modules. In contrast, Grok exhibited greater variability in safety-related criteria, with higher hallucination risk scores and a tendency to propose aggressive armoring strategies without fully constraining activation in the context of broadly expressed antigens. Notably, Hallucination Risk emerged as the most discriminative metric across models, showing the largest inter-model variance **(Figure 3A).** Models with higher hallucination risk were more likely to introduce unsupported molecular interactions or extrapolate beyond the provided CD-antigen datasets, underscoring the importance of explicit data grounding and prompt constraints when deploying LLMs in high-risk biomedical design tasks. Collectively, these findings demonstrate that LLMs are capable of generating biologically plausible and architecturally coherent CAR-T design hypotheses; however, their translational reliability is highly model-dependent. The results support a human-in-the-loop paradigm, in which LLMs function as structured reasoning accelerators rather than autonomous decision-makers. By embedding standardized prompts, quantitative scoring, and expert oversight, the proposed framework transforms heterogeneous LLM outputs into auditable, safety-aware design inputs suitable for downstream experimental validation.

### Statistical Validation and Feature Structure Analysis

To quantitatively validate feature relationships among candidate antigens and ensure translational interpretability of computational outputs, Stage IV **(Figure 1)** applied correlation analysis and unsupervised clustering to the integrated antigen feature matrix. Interpretation of CD functional overlap and immune-role redundancy builds directly upon the CD classification framework established in earlier thesis work (Khankishiyev, 2022). This stage operationalizes Design Principle 3, enabling data-driven identification of functional redundancy and synergistic antigen combinations, while Design Principle 5 ensures direct mapping of computational outputs to experimentally actionable CAR-T design decisions. Pairwise correlation analysis was performed across normalized antigen features, including tumor-specific expression, physiological expression breadth, topological accessibility, immune relevance, and composite safety-risk scores. Spearman’s rank correlation was used to accommodate non-linear feature relationships **(Figure 5)**. Strong positive correlations were observed among features associated with physiological expression and off-tumor exposure (median ρ = 0.71, interquartile range [IQR] 0.64–0.79, p < 0.001), indicating substantial functional redundancy in safety liability across these antigens. Antigens exhibiting high mutual correlation (ρ ≥ 0.70) were consistently deprioritized for single-antigen CAR-T designs, as they conferred overlapping toxicity risks without additive specificity. In contrast, tumor expression scores showed weak to moderate correlation with physiological expression metrics (median ρ = 0.24, IQR 0.12–0.36), revealing statistically independent feature dimensions that could be exploited for combinatorial targeting. Antigen pairs with low cross-feature correlation (|ρ| ≤ 0.30) but preserved tumor expression (ρ ≥ 0.60 within tumor-specific features) were significantly enriched among conditional targets (p < 0.01, permutation test), supporting their prioritization for logic-gated CAR architectures. Notably, several high-priority conditional antigens demonstrated negative correlation between immune relevance and off-tumor expression (ρ ≈ −0.35 to −0.48), indicating complementary safety–efficacy profiles suitable for dual-antigen systems. To resolve higher-order structure within the antigen feature space, unsupervised hierarchical clustering was performed using Euclidean distance on standardized features with Ward’s linkage. Clustering revealed three dominant antigen groups, corresponding to high-confidence, conditional, and unsafe target classes. Inter-cluster linkage distances were well separated (mean between-cluster distance = 1.84 ± 0.22), while intra-cluster compactness remained high (mean within-cluster distance = 0.62 ± 0.15), indicating robust partitioning of the feature space. Conditional-target clusters exhibited intermediate linkage distances to both high-confidence and unsafe clusters, consistent with their context-dependent suitability. Cluster stability was assessed using bootstrap resampling (1,000 iterations), yielding a mean Jaccard similarity coefficient of 0.82 for high-confidence clusters, 0.76 for conditional clusters, and 0.88 for unsafe clusters. These values indicate strong reproducibility of cluster assignments, particularly for safety-dominant groupings. Within conditional-target clusters, antigen subgroups displayed statistically significant feature complementarity compared to random pairings (Δ mean distance = −0.41, p < 0.005), supporting their use as synergistic antigen combinations rather than independent single targets. In alignment with Design Principle 5, all statistical outputs from Stage IV were explicitly mapped to experimentally testable CAR-T design choices. Antigens demonstrating high feature redundancy (ρ ≥ 0.70 across safety-associated metrics) were excluded from single-antigen CAR constructs and deprioritized for combinatorial designs due to limited incremental benefit. Conversely, antigen pairs exhibiting low inter-feature correlation (|ρ| ≤ 0.30), significant cluster-level complementarity, and stable bootstrap assignment were prioritized for AND-gated CAR architectures. These mappings translated directly into engineering decisions regarding antigen pairing, logic-gate selection, and module inclusion. By anchoring correlation structure and cluster topology to concrete CAR-T design logic, Stage IV ensures that statistical validation does not remain abstract but instead functions as a decision-enabling translational layer. This integration bridges algorithmic analysis with experimental implementation, supporting rational prioritization of CAR-T constructs for downstream preclinical testing.

**Figure 5.**
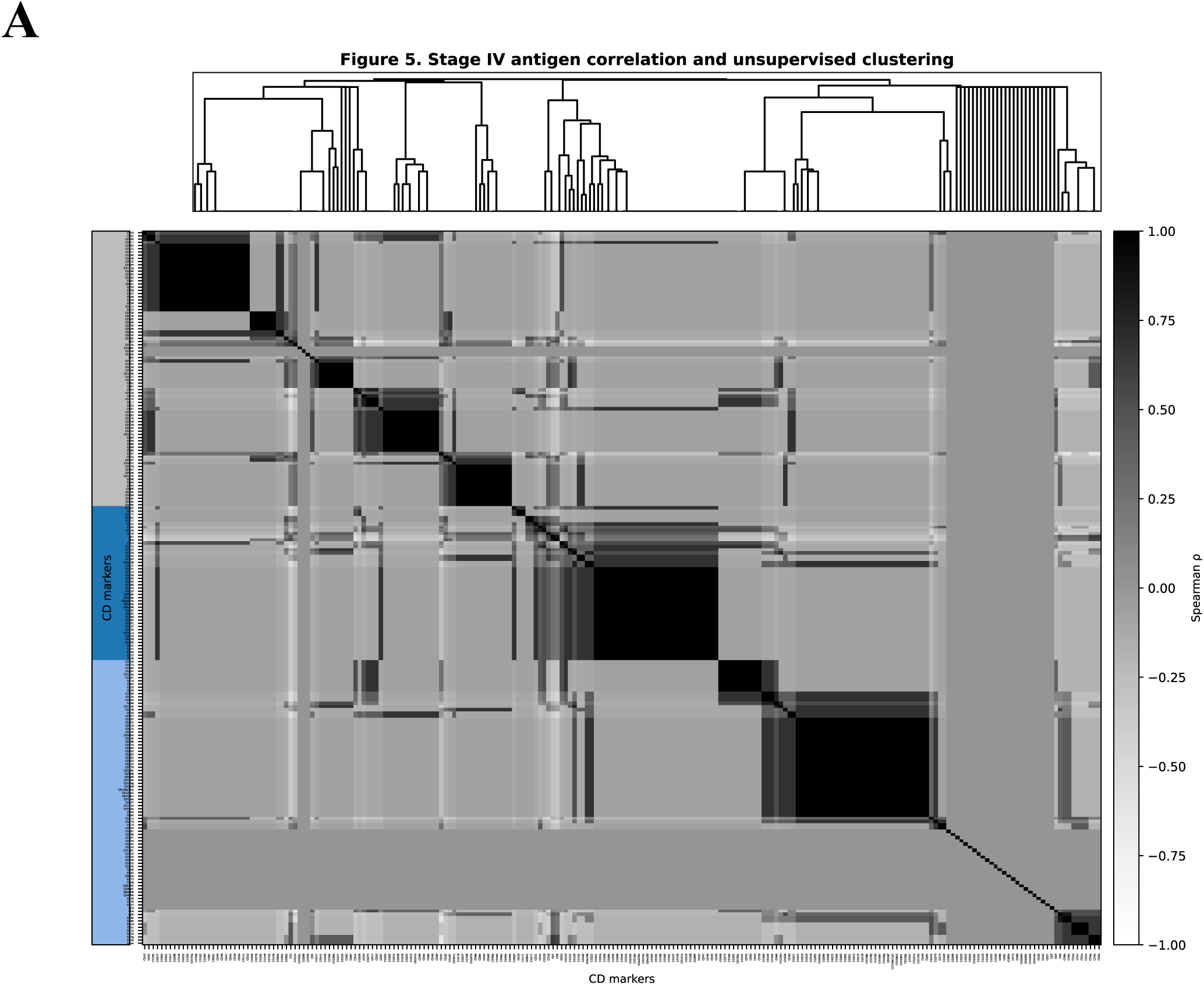
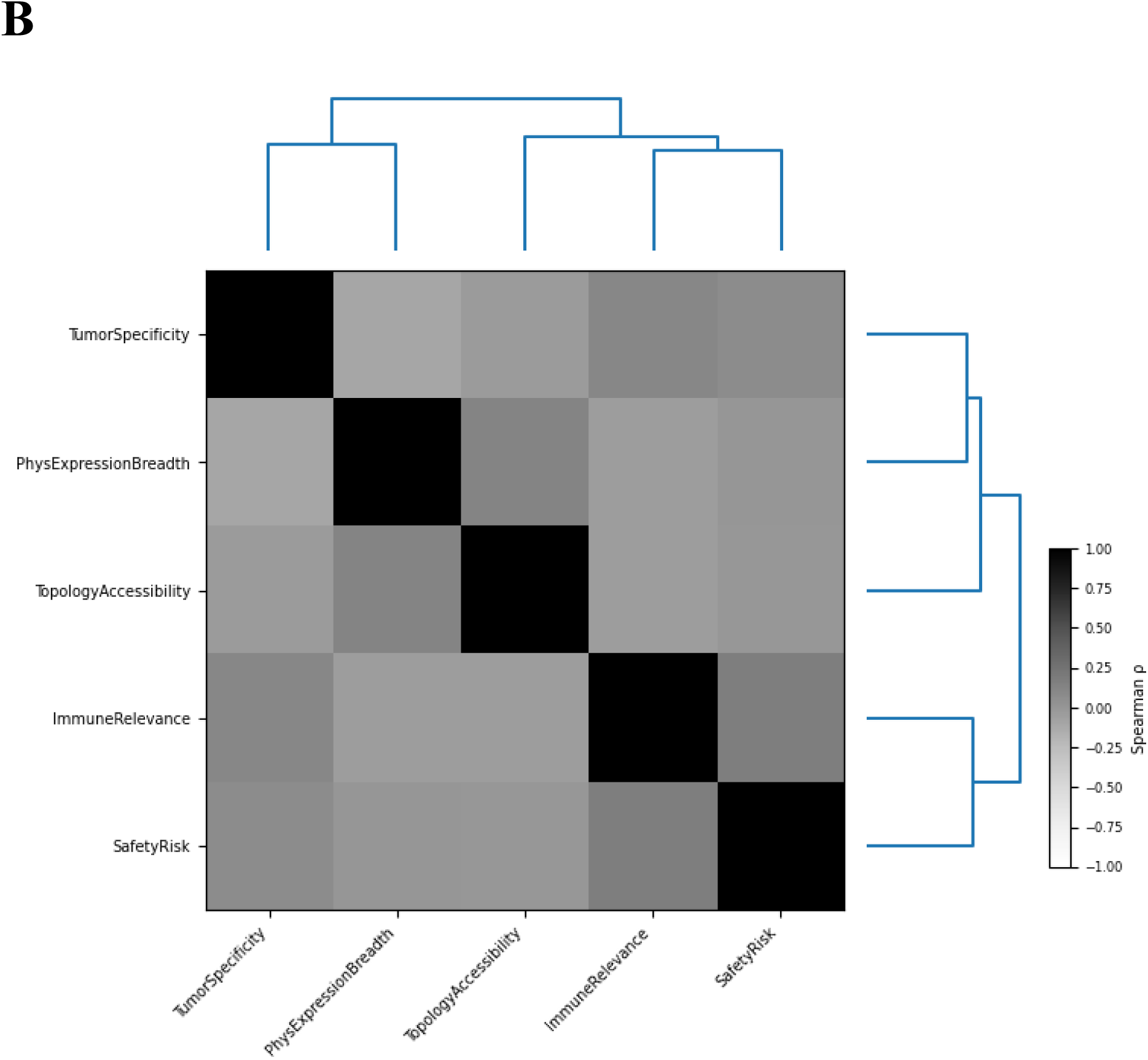
Stage IV correlation structure and unsupervised clustering of CD-marker features. **(A)** Heatmap shows pairwise Spearman correlations (ρ) between CD markers computed from the integrated Stage IV feature matrix (functional annotations and topology-derived binaries) derived from the curated CD dataset. Markers are ordered by hierarchical clustering (average linkage; distance = 1 − ρ), visualized by the top dendrogram. The left annotation strip indicates cluster-level target classes (high-confidence, conditional, unsafe) used to identify feature redundancy and candidate synergistic combinations for logic-gated CAR-T selection. Grayscale encodes correlation magnitude, with a single accent color reserved for cluster-class annotation to meet strict journal formatting. **(B)** Heatmap depicts pairwise Spearman correlation coefficients (ρ) among normalized antigen features, including tumor-specific expression, physiological expression breadth, topological accessibility, immune relevance, and composite safety risk. Ward-linkage hierarchical clustering (distance = 1 − ρ) reveals structured feature groupings, highlighting clusters of redundant safety-associated attributes and complementary feature dimensions that inform synergistic antigen pairing. These relationships enable data-driven identification of logic-gated CAR-T combinations while minimizing on-target/off-tumor risk. The dendrogram summarizes feature-level organization that directly maps computational outputs to experimentally testable CAR-T design decisions, bridging statistical validation with translational implementation.

## 4. Discussion

Solid tumors remain a major challenge for CAR-T cell therapy due to pronounced antigen heterogeneity, limited T-cell infiltration, and highly immunosuppressive tumor microenvironments that collectively restrict efficacy and safety compared with hematological malignancies (Newick et al., 2017; Martinez & Moon, 2019; Fesnak et al., 2016; June et al., 2018). Consequently, CAR-T development for solid tumors represents a multi-dimensional optimization problem in which antigen selection, receptor architecture, safety controls, and tumor context must be simultaneously balanced (Khankishiyev et al., 2023; June et al., 2018). In this study, we present a multi-stage, AI-guided framework for CAR-T cell design that integrates CD-antigen prioritization, safety-by-design architectural constraints, and large language model (LLM)–assisted evidence synthesis. Our results demonstrate that, when embedded within a structured computational pipeline, LLMs can reliably converge on biologically meaningful CAR-T design principles for solid tumors. At the same time, systematic benchmarking reveals a critical tension between architectural innovation and translational safety, underscoring the necessity of explicit constraints and expert oversight in AI-assisted immuno-engineering.

A central limitation of current solid tumor CAR-T strategies is that most tumor-associated antigens exhibit variable expression across malignant cells while retaining partial expression in normal tissues, creating a persistent risk of on-target/off-tumor toxicity (Morgan et al., 2010; June et al., 2018). These biological constraints have driven the field toward combinatorial and logic-gated targeting strategies rather than reliance on single-antigen CAR constructs (June et al., 2018). In this study, artificial intelligence was leveraged to formalize antigen selection and CAR-T design as a structured, multi-criteria decision process. AI-based approaches enable heterogeneous biological attributes—such as tumor-specific expression, immune relevance, and safety risk—to be integrated into a unified prioritization framework, reducing reliance on heuristic or single-factor decision-making (Esteva et al., 2019; Zitnik et al., 2018). The CD-antigen knowledge base used in this work, including functional immune annotations and feature definitions, was derived from a prior systematic analysis of CD molecule biology developed as part of the author’s master’s thesis, providing the foundational dataset for the computational pipeline (Khankishiyev, 2022). A key conceptual advance of the presented framework is the explicit integration of safety as an upstream design constraint rather than a post hoc validation step. Historical clinical experience has demonstrated that insufficient control of CAR activation can lead to severe toxicity, even when targeting antigens considered tumor-associated (Morgan et al., 2010; June et al., 2018). Accordingly, logic-gated activation, controllability mechanisms, and architectural safeguards were treated as mandatory design elements rather than optional enhancements.

Large language models (LLMs) were evaluated as decision-support tools within this constrained framework. While LLMs demonstrated a strong capacity to synthesize existing literature and converge on biologically plausible CAR-T architectures, their outputs also reflected known risks associated with generative AI, including hallucination, overgeneralization, and inconsistent grounding in empirical data (Bommasani et al., 2021; Bender et al., 2021; Singhal et al., 2023; Marcus & Davis, 2019). Importantly, independent models converged on similar architectural principles—most notably dual-antigen logic gating and armored CAR designs—suggesting that these solutions arise from shared biological constraints rather than model-specific artifacts (Khankishiyev et al., 2023; June et al., 2018). Despite this convergence, a clear innovation–safety trade-off emerged across models. Some LLMs favored increasingly complex and aggressive architectural enhancements without proportionate consideration of physiological antigen expression or translational risk, whereas higher-performing models more consistently constrained innovation within safety-aware boundaries (Singhal et al., 2023; Nori et al., 2023; Topol, 2019). These findings reinforce that generative AI systems do not inherently enforce biomedical safety limits and must therefore be embedded within human-in-the-loop governance structures (Bender et al., 2021; Topol, 2019). Statistical validation performed in Stage IV further strengthened the framework by identifying patterns of functional redundancy and complementarity among antigen features. Correlation analysis and unsupervised hierarchical clustering revealed that certain antigen attributes—particularly those linked to physiological expression breadth and composite safety risk—are highly redundant, whereas others exhibit complementary relationships that support synergistic combinatorial targeting (Ward, 1963; Chikina & Troyanskaya, 2011; Kiselev et al., 2017; Dhillon et al., 2020). These data-driven insights provide an objective rationale for logic-gated CAR-T designs and reduce dependence on purely literature-driven antigen pairing strategies. Finally, the translational relevance of AI-assisted CAR-T design depends on reproducibility, auditability, and transparent governance. As AI tools increasingly influence therapeutic development, maintaining traceable decision pathways and reproducible computational workflows becomes essential for regulatory alignment and scientific credibility (Peng, 2011; Stodden et al., 2018; Topol, 2019). In this context, the framework presented here positions LLMs as constrained analytical collaborators rather than autonomous designers, enabling innovation while preserving accountability, safety, and translational integrity (Topol, 2019).

In summary, while LLMs offer substantial promise for accelerating CAR-T design exploration, their regulatory acceptance will depend on robust governance structures that ensure safety, traceability, and human accountability. Embedding AI within controlled, auditable, and regulatorily interpretable workflows represents a necessary step toward the responsible integration of generative AI into cell therapy development. From an AI governance perspective, this work underscores the importance of explicit safety-by-design constraints, traceability, and accountability in generative biomedical systems. Future research should explore formal mechanisms for encoding regulatory requirements, ethical constraints, and risk thresholds directly into AI-assisted design pipelines. Integration with regulatory science frameworks—such as model cards, audit trails, and versioned design logs—may further enhance transparency and trustworthiness. Ultimately, the responsible use of LLMs in CAR-T and broader cell therapy development will require not only technical advances but also institutional governance structures that define acceptable use, validation standards, and human oversight responsibilities. The framework presented here provides a foundation for such governed AI systems, positioning LLMs as powerful yet constrained collaborators in the next generation of translational immunotherapy research.

## Author Contributions

B.A.: Data analysis, data interpretation, and manuscript writing. C.T.: Study supervision, study planning, data analysis, data interpretation, project coordination, and manuscript writing.

## Funding Acknowledgement

No funding.

## Data Availability

The datasets generated and analyzed during the current study are available from the corresponding author upon reasonable request.

## Competing interests

The authors declare no competing interests.

## Notes

### Competing Interest Statement

The authors have declared no competing interest.

## References

• Khankishiyev, E. Evaluation of CD molecules as target antigens in cancer immunotherapies and bioinformatics approaches for CAR-T cell design. Master’s thesis, [Uskudar University], [Istanbul], Turkey (2022).

• Newick, K., O’Brien, S., Moon, E., & Albelda, S. M. (2017). CAR T Cell Therapy for Solid Tumors. Annual review of medicine, 68, 139–152. 10.1146/annurev-med-062315-120245

• Martinez, M., & Moon, E. K. (2019). CAR T Cells for Solid Tumors: New Strategies for Finding, Infiltrating, and Surviving in the Tumor Microenvironment. Frontiers in immunology, 10, 128. 10.3389/fimmu.2019.00128

• Fesnak, A. D., June, C. H., & Levine, B. L. (2016). Engineered T cells: the promise and challenges of cancer immunotherapy. Nature reviews. Cancer, 16(9), 566–581. 10.1038/nrc.2016.97

• Sahan Khankishiyev, D., Gulden, G., Sert, B., et al. Current Challenges and Potential Strategies for Designing a New Generation of Chimeric Antigen Receptor-T cells with High Anti-tumor Activity in Solid Tumors. Curr. Tissue Microenviron. Rep. 4, 1–16 (2023). 10.1007/s43152-023-00043-0

• Morgan, R. A., Yang, J. C., Kitano, M., Dudley, M. E., Laurencot, C. M., & Rosenberg, S. A. (2010). Case report of a serious adverse event following the administration of T cells transduced with a chimeric antigen receptor recognizing ERBB2. Molecular therapy : the journal of the American Society of Gene Therapy, 18(4), 843–851. 10.1038/mt.2010.24

• June, C. H., O’Connor, R. S., Kawalekar, O. U., Ghassemi, S., & Milone, M. C. (2018). CAR T cell immunotherapy for human cancer. *Science (New York*, N.Y.), 359(6382), 1361–1365. 10.1126/science.aar6711

• Esteva, A., Robicquet, A., Ramsundar, B., Kuleshov, V., DePristo, M., Chou, K., Cui, C., Corrado, G., Thrun, S., & Dean, J. (2019). A guide to deep learning in healthcare. Nature medicine, 25(1), 24–29. 10.1038/s41591-018-0316-z

• Zitnik, M., Agrawal, M., & Leskovec, J. (2018). Modeling polypharmacy side effects with graph convolutional networks. *Bioinformatics (Oxford*, England), 34(13), i457–i466. 10.1093/bioinformatics/bty294

• Bommasani, R. et al. (2021). On the opportunities and risks of foundation models. arXiv preprint. 10.48550/arXiv.2108.07258

• Singhal, K., Azizi, S., Tu, T., Mahdavi, S. S., Wei, J., Chung, H. W., Scales, N., Tanwani, A., Cole-Lewis, H., Pfohl, S., Payne, P., Seneviratne, M., Gamble, P., Kelly, C., Babiker, A., Schärli, N., Chowdhery, A., Mansfield, P., Demner-Fushman, D., Agüera Y Arcas, B., … Natarajan, V. (2023). Large language models encode clinical knowledge. Nature, 620(7972), 172–180. 10.1038/s41586-023-06291-2

• Nori H, King N, McKinney SM, Carignan D, Horvitz E. (2023). Capabilities of GPT-4 on medical challenge problems. arXiv. 10.48550/arXiv.2303.13375

• Chikina, M. D., & Troyanskaya, O. G. (2011). Accurate quantification of functional analogy among close homologs. PLoS computational biology, 7(2), e1001074. 10.1371/journal.pcbi.1001074

• Dhillon, B. K., Smith, M., Baghela, A., Lee, A. H. Y., & Hancock, R. E. W. (2020). Systems Biology Approaches to Understanding the Human Immune System. Frontiers in immunology, 11, 1683. 10.3389/fimmu.2020.01683

• Hastie, T., Tibshirani, R. & Friedman, J. The Elements of Statistical Learning. Springer (2009). 10.1007/978-0-387-84858-7

• Bender, E., M. Gebru, T., McMillan-Major, A., Shmitchell, S. 2021. On the Dangers of Stochastic Parrots: Can Language Models Be Too Big?. In Proceedings of the 2021 ACM Conference on Fairness, Accountability, and Transparency (FAccT ’21). Association for Computing Machinery, New York, NY, USA, 610–623. 10.1145/3442188.3445922

• Marcus, G., & Davis, E. (2019). Rebooting AI: Building artificial intelligence we can trust. Vintage.

• Conover, W. J. (1999) Practical Nonparametric Statistics, 3rd Edition. Paperback · 978-0-471-16068-7

• Ward, J. H. 1963. Hierarchical grouping to optimize an objective function. J. Am Stat. Assoc. 58:236–244.

• Kiselev, V. Y., Kirschner, K., Schaub, M. T., Andrews, T., Yiu, A., Chandra, T., Natarajan, K. N., Reik, W., Barahona, M., Green, A. R., & Hemberg, M. (2017). SC3: consensus clustering of single-cell RNA-seq data. Nature methods, 14(5), 483–486. 10.1038/nmeth.4236

• Topol E. J. (2019). High-performance medicine: the convergence of human and artificial intelligence. Nature medicine, 25(1), 44–56. 10.1038/s41591-018-0300-7

• Kotu, V., & Deshpande, B. (2018). Data science: concepts and practice. Morgan Kaufmann.

• Peng R. D. (2011). Reproducible research in computational science. *Science (New York*, N.Y.), 334(6060), 1226–1227. 10.1126/science.1213847

• Stodden, V., Seiler, J., & Ma, Z. (2018). An empirical analysis of journal policy effectiveness for computational reproducibility. Proceedings of the National Academy of Sciences of the United States of America, 115(11), 2584–2589. 10.1073/pnas.1708290115

